# Liposomal aggregates sustain the release of rapamycin and protect cartilage from friction

**DOI:** 10.1101/2023.03.23.533793

**Authors:** Gregor Bordon, Shivaprakash N. Ramakrishna, Sam G. Edalat, Remo Eugster, Andrea Arcifa, Simone Aleandri, Mojca Frank Bertoncelj, Lucio Isa, Rowena Crockett, Oliver Distler, Paola Luciani

## Abstract

Fibrosis, low-grade inflammation, and increased friction are together with degradation of cartilage key culprits for debilitating pain in osteoarthritis (OA), which is one of the most common chronic diseases of today’s aging population. Intraarticular administration of bio-lubricants loaded with a pharmaceutically active component recently showed promise to improve therapy. Liposomes have emerged as exceptional lubricant biomaterial, but their small size leads to rapid clearance from the synovium, causing a need for more frequent administration. We recently developed a liposomal drug delivery system based on aggregation of negatively charged liposomes with physiologically present divalent cations. Here, we expanded our platform by replacing calcium with zinc, reported to exert anti-inflammatory action. The liposomal aggregates extend the release of rapamycin (RAPA) beyond the free liposomes and have a diameter of nearly 100 μm, which was previously established to improve retention in synovial joints. Electron microscopy showed that RAPA alters the irregular morphology of liposomal clusters, which are irreversible upon dilution. RAPA recently showed great promise both *in vitro* and *in vivo* at protecting the joints from inflammation and cartilage from further degradation. Our study adds to this by showing that RAPA is also able to dampen the fibrotic response in human OA synovial fibroblasts. Finally, the tribological properties were assessed on nano- and macro-scales on silicon surface and *ex vivo* porcine cartilage, which showed an excellent protective ability of the system against friction on both scales. Taken together, our study shows that liposomal aggregates have the potential of improving local OA therapy.

## 1 Introduction

Osteoarthritis (OA) is a debilitating chronic joint disease characterised by the degradation of articular cartilage, synovial fibrosis, and low-grade inflammation. It affects 7% of the global population, where women are disproportionately affected by the condition. The current treatment possibilities are very limited, relying primarily on non-steroidal anti-inflammatory drugs and analgesics, and joint replacement. For patients unresponsive to these medications, hyaluronic acid (HA) and glucocorticoids are prescribed, however, their use remains controversial. Several cellular therapies are becoming increasingly available, but they lack consistency of protocols or/and strong clinical data [1–3]. Overall, better treatments for OA remain a strongly unmet clinical need.

Rapamycin (RAPA) is an immunosuppressive drug that was first approved for the prevention of rejection in renal transplant recipients [4] and has been since tested for treatment of cancer [5], inflammatory diseases [6–8], and increasing longevity [9,10]. Recently, RAPA showed promise in OA therapy, as it reduces excessive chondrocyte apoptosis and inflammation, protecting the cartilage from further degradation [11,12]. Several in vivo studies have confirmed these effects, demonstrating that RAPA significantly reduces OA severity and damage to the articular cartilage [13–15]. The latest evidence shows that OA also perturbs the function of synovial fibroblasts (SFs) in the joint synovial membrane. SFs significantly contribute to cartilage damage in OA, and synovial fibrosis is associated with chronic joint pain [2,16,17]. While research on RAPA’s effects on SF inflammation and senescence is gathering momentum [18–20], little is known about its impact on fibrotic OA SFs (OASFs). Additionally, the systemic use of RAPA is hindered by its adverse effects, but a growing body of literature suggests that local intraarticular injection is a promising avenue for OA treatment [13,14].

Previous research has proposed that a combination of pharmacological intervention and cartilage lubrication would yield a synergistically improved treatment for OA, but there are still no therapies available for patients exerting this dual activity [21,22]. Early investigations demonstrated that phospholipid-based liposomes, namely small unilamellar vesicles (SUVs), can improve boundary lubrication and wear of cartilage through hydration of the phospholipid headgroups [23–26]. More recently, scientists developed a liposomal system that sustained the drug release of D-glucosamine sulfate for OA treatment and succeeded in improving the lubrication [22]. Although liposomes show great promise for OA treatment from the standpoint of biocompatibility, boundary lubrication, and controlled drug release, typical SUVs suffer from several drawbacks. Small SUVs can penetrate deep into cartilage and are on more physiological models worse lubricants than larger phospholipid-based vesicles that can be retained closer to the tissue’s surface [27]. Furthermore, a small particle size of below 300 nm was correlated with rapid clearance from the joint, which calls for frequent administration of the formulation and an increased risk of inducing infection [28,29]. In comparison, particles above 10 μm can avoid phagocytosis by macrophages and can be retained in naïve as well as in the inflamed joints for over 6 weeks [29–31]. For these reasons, the utility of small phospholipid-based particles, such as SUVs is in practice limited and the development of new drug delivery systems with larger particle size and ability to sustain the drug release is imperative for better treatment of OA. Our group previously reported the ability of calcium and magnesium cations to aggregate the negatively charged liposomes into larger aggregated liposomes (ALs) forming injectable depots [32] and the resulting slower release of bupivacaine *in vitro* [33]. *In vivo* results suggested that the aggregates also increased the drug’s area under the curve in plasma compared to the non-depot system and modulated the particle clearance from the injection site. Here, we tested zinc, a divalent cation known for its anti-inflammatory and antioxidant effects [34–36], as an alternative to the aforementioned aggregating agents. Our results demonstrate that aggregating negatively charged liposomes with 150 mM zinc produces irreversible ALs (ZnALs) with a diameter exceeding 90 μm, which has been shown to significantly increase the retention time in synovial joints. [29,31]. The irreversible nature of the particles is not significant only from a pharmacokinetic perspective, but also technological, because it allows downstream processing, such as purification from excess zinc prior to administration. We further characterised the aggregation properties of the system in depth and showed that ZnALs are able to improve lubrication through testing with lateral force microscopy as well as sustain the release of RAPA with 86 % of the drug released after 7 days. These findings suggest that the system has potential for the dual treatment of OA through maintaining a low level of friction in the joint and sustaining the release of RAPA, which can decrease fibrotic markers in OASFs.

## 2 Materials and methods

### 2.1 Materials

The phospholipids 1,2-dipalmitoyl-*sn*-glycero-3-phosphocholine (DPPC) and 1,2-distearoyl-*sn*-glycero-3-phospho-(10-rac-glycerol) sodium salt (DSPG) were kindly gifted by Lipoid (Ludwigshafen, Germany). Rapamycin (sirolimus) was obtained from R&S Pharmchem (Pudong Districs, Shanghai, China). Zinc chloride (98% purity, reagent grade), ketoconazole (99-101% purity), and cholesterol (≥99% purity) were purchased from Sigma-Aldrich-Merck (St Louis, MO, USA). Trifluoroacetic acid and 1 M HEPES solution were obtained from Carl Roth (Karlsruhe, Germany). 1,1’-dioctadecyl-3,3,3’,3’-tetramethylindodicarbocyanine, 4-chlorobenzenesulfonate salt (DiD, catalog number: D7757) was purchased from Thermo Fisher Scientific (Waltham, MA, USA). Chloroform and methanol were obtained from Fisher Scientific (Schwerte, Germany). All chemicals were used as received. Ultrapure water of resistivity 18.2 MΩ.cm was produced by a Barnstead Smart2 pure device from Thermo Scientific (Pittsburgh, USA). Porcine knee cartilage was obtained from a local slaughterhouse in Münchenbuchsee, Switzerland.

### 2.2 Preparation and characterisation of liposomes

Liposomes with 25 mol% DSPG and varying DPPC/cholesterol content were prepared with thin-film hydration method. Lipid stock solutions in a chloroform/MeOH mixture (75/25 v/v) were dried under nitrogen flow and kept under vacuum overnight to remove residual solvents. Vesicles of 20 mM final lipid concentration were formed by hydration with 20 mM HEPES buffer at pH 7.4, heating to 70 °C, and mixing. The formed vesicles were freeze-thawed 6 times and subsequently extruded 10 times through a 200 nm polycarbonate membrane (Sterlitech Corporation, USA) with a LIPEX extruder at 70 °C (Evonik, Canada). The mean hydrodynamic diameter and polydispersity index (PDI) were measured with dynamic light scattering (DLS) analyser Litesizer 500 (Anton Paar, Austria) at 25 °C with a backscatter angle of 175° and a 658 nm laser. The zeta potential was assessed with laser Doppler microelectrophoresis using the same instrument and Omega cuvette (Anton Paar, Austria). Liposome stability was evaluated for 16 weeks at 4 °C. Formulations were used within 24 h from extrusion for all testing.

### 2.3 Encapsulation efficiency of RAPA

RAPA was dissolved in MeOH and added to the lipid film in varying molar lipid: drug ratios, by keeping the lipid content constant. Encapsulation efficiency was calculated according to the formula below:

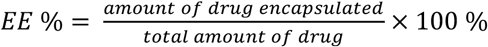

The encapsulated RAPA was quantified after removal of unencapsulated RAPA with preparative size exclusion chromatography (SEC) column (PD MidiTrap, G-25, Cytiva, USA) according to the manufacturer’s protocol. The drug concentration in samples was measured with high-performance liquid chromatography (HPLC) using a reverse phase C18 Nucleosil 100-5 (4.0 × 250 mm; 5.0 μm particle size, Macherey-Nagel, Germany) column and mobile phase consisting of MeOH/water (90/10 v/v) + 0.1% trifluoroacetic acid at a flow rate of 1 mL/min, temperature 50° C and UV detection at λ = 278 nm. Ketoconazole was added to all samples as an internal standard at concentration of 0.2 mg/mL.

### 2.4 *In vitro* drug release

Release of RAPA from liposomes and ZnALs was tested *in vitro* with a custom-made dialysis device (**Figure S1**), with the same dimensions as a 2 mL Slide-A-Lyzer MINI (Thermo Scientific, USA) and a disposable polycarbonate membrane with 100 nm pore size (Figure S1). The release medium contained 10% EtOH in ultrapure water to achieve sink conditions and RAPA stability. A volume of 1 mL of each sample was added into the dialysis device and 48 mL of the release medium were added to the acceptor chamber. The loaded dialysis devices were placed in the incubator at 37 °C while shaking at 10 rpm. Throughout the 7 days of the study’s duration, the release medium from the acceptor chamber was aliquoted and fully replaced with a fresh one at each time point. The aliquots were frozen in liquid nitrogen and lyophilised. Each sample was resuspended with the internal standard solution and RAPA content was determined with HPLC.

### 2.5 Preparation and characterisation of ZnALs

Aggregated liposomes (ALs) were prepared through a four-time dilution of 20 mM liposomes with aqueous solutions containing different concentrations of ZnCl2 (Zn^2+^) and gentle stirring for 5 min. ALs were characterised via turbidimetric scattering measurements using a microplate reader, as previously reported [32]. Briefly, 50 μL of liposomes with an initial lipid concentration of 20 mM were mixed with 150 μL Zn^2+^ solution in a quartz 96-well microtiter plate with a clear and flat bottom (Hellma GmbH & Co. KG, Germany). The mixture was gently mixed for 30 min, and the optical density was measured with an Infinite M Pro 200F-PlexNano microplate reader (Tecan, Switzerland) at 450 nm. The zeta potential of ZnALs was measured with the Litesizer 500 in the same manner as for the liposomes. Stability upon dilution was tested by diluting ALs 400 times with ultrapure water and subsequently performing a size measurement with the

DLS at different time points. The presence and size of aggregates were determined with laser diffraction measurements, which were performed with PSA 1190 LD (Anton Paar GmbH, Austria) after 400x dilution in water as a dispersion medium where the obscuration parameter was set to 1-7 %. The optimization of input parameters such as stirring and pump speed was defined in order to obtain repeatable measurements. Stirring was set to slow (150 rpm) and the pump speed was put to medium setting (120 rpm). To further confirm the presence of aggregates upon 400x dilution, a nanoparticle tracking analysis (NTA) was performed using Zetaview (Particle Metrix, Germany) with a 488 nm laser, camera sensitivity of 58 and a 100 m/s shutter value. Aggregation kinetic experiments were performed with PSA using a series of 180 measurements with a measurement time of 10 s between each point. D50 was recorded and plotted over time, where 3 time points were averaged together as technical repeats to account for the measurement fluctuations. Prior to testing, the ZnALs underwent a purification process to eliminate excess Zn^2+^. This was achieved through a 4 h dialysis in 0.5 L of ultrapure water, utilizing Float-A-Lyzers G2, 8-10 kDa MWCO (LubioScience GmbH, Switzerland), and hourly complete replacement of medium. Zn^2+^ concentration was measured using inductively coupled plasma mass spectrometry (ICP-MS, NexION 2000, PerkinElmer, USA). Calibration standards, internal standards, and samples were prepared using a 2% (w/w) HNO3 solution (BASF SE, Germany) as the matrix. During measurements, a 10 μg/L yttrium solution was utilized as the internal standard during measurements. The ICP-MS system was calibrated with standards (TraceCERT Merck, Germany) ranging from 1 to 500 μg/L Zn^2+^.

### 2.6 Differential scanning calorimetry (DSC)

Interactions between the phospholipid bilayer and RAPA were examined with a DSC 250 (TA Instruments, USA). Multilamellar liposomes and ZnALs (150 mM Zn^2+^) with or without RAPA were prepared in a final concentration of 20 mM and 12 μL were transferred in a Tzero^®^ aluminum pan and hermetically sealed. A volume of 12 μL of 20 mM HEPES was used as a reference. Samples were pre-heated to 60 °C, kept at that temperature for 5 min, and then cooled down to 10 °C. Next, two heating and cooling cycles were performed between 10 °C and 70 °C at the rate of 2 °C/min. The last cycle was used for the evaluation of the thermal profile and the calculation of hysteresis. The enthalpy values were normalized to the phospholipid amounts in the samples.

### 2.7 Microscopic imaging of liposomes and ZnALs

Liposomes and ZnALs were imaged with fluorescence and cryogenic transmission electron microscopy (cryo-TEM) to assess the morphology. For fluorescence microscopy, lipid films for liposome preparation were stained with 0.05 mol% of the non-exchangeable lipophilic dye DiD. Liposomes and ZnALs were prepared as described above, while being protected from light. A volume of 20 μL of formulations was added on a slide and covered with a glass coverslip to be imaged with an inverted fluorescence microscope (Nikon Eclipse-Ti, Canada) through Tx red filter. For cryo-TEM, a volume of 6-8 μL of each sample was applied onto a gold grid covered by a holey gold film (UltrAuFoil 2/1, Quantifoil Micro Tools GmbH, Jena, Germany). Excess of liquid was blotted automatically between two strips of filter paper or only from the backside of the Grid. Subsequently, the samples were rapidly plunge-frozen in liquid ethane (cooled to 180 °C) in a Cryobox (Carl Zeiss NTS GmbH, Oberkochen, Germany). Excess ethane was removed with a piece of filter paper. The samples were transferred immediately with a Gatan 626 cryo-transfer holder (Gatan, Pleasanton, USA) into the pre-cooled Cryo-electron microscope (Philips CM 120, Eindhoven, Netherlands) operated at 120 kV and viewed under low dose conditions. The images were recorded with a 2k CMOS Camera (F216, TVIPS, Gauting, Germany). In order to minimize the noise, four images were recorded and averaged to one image.

### 2.8 Cell culture

SFs were obtained from four consenting OA patients (according to ethics approvals BASEC-Nr. 2019-00674 and BASEC Nr. 2019-00115) and plated onto 25 cm^2^ flasks and 6-well (clear, Corning, USA) or 96-well (black with clear bottom, Thermo Fisher Scientific, USA) plates following standard protocols [37]. Cells were cultured at 37 °C in a humidified atmosphere at 5% CO2 with Dulbecco’s modified Eagle’s medium (DMEM; Life Technologies) supplemented with 10% fetal calf serum (FCS), 50 U mL^−1^ penicillin/streptomycin, 2 mM L-glutamine, 10 mM HEPES, and 0.2% amphotericin B (all from Life Technologies). OASFs were used for experiments between passages 4 and 6 when they reached confluency.

### 2.9 Toxicity of RAPA and ZnALs

OASFs were counted with Countess 3 FL (Thermo Fisher Scientific, USA) using trypan blue and seeded onto a 96-well plate at a density of 5’000 cells per well. After overnight incubation, the cells were treated with 200 μL of different conditions and incubated for 48 h. Upon incubation, the cells were stained with the LIVE/DEAD™ Viability/Cytotoxicity Assay Kit (Thermo Fisher Scientific, USA) according to the manufacturer’s protocol. Briefly, the medium was aspirated, and the cells were washed with DPBS (Thermo Fisher Scientific, USA) before being stained with calcein and Sytox Deep Red dyes. Cells were incubated for 30min at RT and washed with DPBS. The dead control was prepared by fixing the cells in ice-cold ethanol as per manufacturer’s recommendations. Fluorescence was measured with a plate reader (BioTek Instruments, USA) in a bottom area scan mode with a 35 gain. Fluorescence intensity of alive cells was measured with a 528/20 nm filter after the excitation at 485 nm wavelength. Dead cells’ fluorescence was excited at 530 nm and the emission was detected with 590/35 nm filter. Viability percentage was normalised to the fluorescence intensity of the untreated condition, which was treated with normal medium. Images were taken with a widefield fluorescence microscope (Zeiss AxioObserver Z1, Germany) using GFP (cyan) and DsRed (green) fluorescence filters.

### 2.10 Gene expression

For gene expression experiments, cells were seeded in the same way as described for the toxicity experiments above. The fibrotic response was stimulated by adding 10 ng/mL of TGFβ to the medium together with the tested conditions. After 48 h incubation, cells were lysed, and RNA was extracted with a Quick-RNA Microprep Kit (Zymo Research, USA) as well as on-column DNase I digested according to the manufacturer’s protocol. The purity and amount of RNA were determined by measuring the OD at a ratio of 260 to 280 nm with Nanodrop (Thermo Fisher Scientific). RNA was reverse transcribed and SYBRgreen real-time PCR was performed. Data were analysed with the comparative CT methods and presented as 2^−ΔΔCT^ (i.e., x-fold) as described elsewhere [38] using RPLP0 as a housekeeping gene for sample normalization. Primer sequences are available in Supplementary Information.

### 2.11 Nanotribology

Nanotribology measurements were performed with colloidal probe lateral force microscopy (CP-LFM) using a Bruker Dimension Icon AFM with tipless Au-coated cantilevers (CSC-38, Mikromash, Bulgaria). Cantilever spring constants were determined using the thermal-noise method [39] for normal spring constants and Sader’s method [40] for torsional spring constants. Approximately 8 μm diameter silica particles (EKA Chemicals AB, Kromasil R) were attached to the end of the cantilever using two-component epoxy glue via a home-built micromanipulator. Four different colloidal probes were prepared and treated with UV/ozone for 30 min before measurement. Four silicon wafers (~1×1 cm) were treated with UV/ozone and used as substrates. Prior to each measurement, substrates were immersed in either PBS, Zn^2+^ solution, ZnAL, or liposome solutions (5 mM) for 30 min and rinsed with ultrapure water [22]. CP-LFM measurements were conducted in buffer solution, and friction loops were recorded by scanning the cantilever laterally over the surface. For each applied load, at least five friction loops were acquired, and the average friction values were obtained from trace and retrace curves. Coefficient of friction (COF) values were determined from friction force vs normal force graphs using Amontons’ law (F=μL), where F is friction force, μ is the COF, and L is the normal force. Lateral-force calibration was conducted using the “test-probe method” described by Cannara et al [41] by moving a test probe (a reference cantilever glued with a silica particle of diameter ~40 μm) laterally into contact with a silicon wafer (1×1 cm) used as a “hard wall” to obtain lateral sensitivity values.

### 2.12 Macrotribology

The macroscopic friction behaviour of self-mated cartilage sliding in the presence of measured samples was investigated with a UMT-2 tribometer (Bruker, USA) operating in linear reciprocating mode. The counterparts consisted of two portions of cartilage that were cut from porcine knee and stored at −20 °C until use. Cartilage was glued to the upper and lower parts of the tribometer, respectively, shortly before testing. During each experiment, the upper specimen (5×5 mm) was pressed with a load of 1 N against the lower specimen (20×10 mm) and rubbed over a 2 mm stroke length at a frequency of 1 Hz for 10min. The pair was completely immersed in the lubricant for the entire duration of the test. All tests were conducted at a constant temperature of 20°C and a data acquisition frequency of 500 Hz. The representative coefficient of friction (COF) of each test was computed from the raw data of lateral and normal force as the average of each friction loop, taking only the central 90% portion of each friction loop to avoid transients associated with the two ends of the stroke length. In addition, the first 20% of the loops were excluded to consider the steady-state friction behaviour of the self-mated contact, thus excluding possible running-in transients.

### 2.13 Statistical analysis

All experiments were carried out in at least three replicates unless otherwise stated. The reported values are means with ± standard deviation. Microsoft Excel was used for general calculations, while GraphPad Prism 9.5 was used for plotting, performing the one-way ANOVA and Tukey’s test.

## 3 Results & Discussion

### 3.1 Preparation of liposomes and drug encapsulation

Drugs with low water solubility, such as RAPA, must be formulated to increase their bioavailability. Encapsulating them in liposomes is an effective way to solubilize the molecules, enhance their therapeutic index, and enable controlled release [21,42]. A key factor in liposome production is the encapsulation efficiency (EE%), with a low EE% requiring further purification to remove the unencapsulated drug, leading to increased production cost and complexity. The EE% is highly dependent on the drug’s chemical properties, the composition of the liposomes, and the lipid-to-drug ratio (L/D) [43]. The impact of both was investigated in **Figure 1)**, where the higher L/D yielded higher EE%.

**Figure 1:**
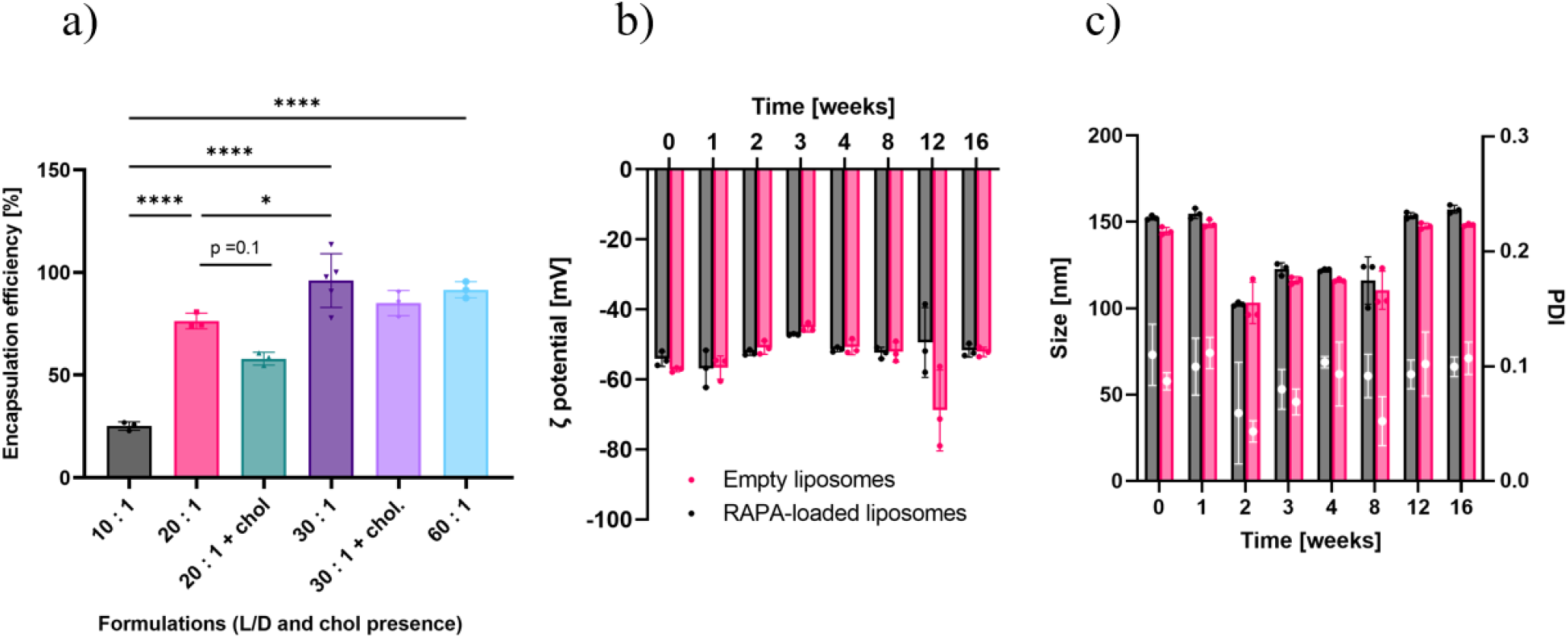
a) EE% of RAPA in liposomes at different L/D. The formulations without chol had the composition DPPC:DSPG = 75:25 and the formulations with chol DPPC:DSPG:chol = 45:25:30. b) Stability of zeta potential over 16 weeks. c) Stability of liposomes over 16 weeks with size on the left y-axis represented in bars and PDI on the right y-axis represented with the scatter plot. One-way ANOVA with Tukey’s multiple comparisons test was run. Statistical significance is designated as: *P < 0.05 **P < 0.01, ***P < 0.001, ****P < 0.0001.

This is to be expected, as RAPA is highly hydrophobic and a greater amount of phospholipid gives more space for the drug to be encapsulated within the bilayer [44]. Chol is oftentimes added to the formulations to modulate the membrane properties such as thickness, packaging, and fluidity, which in turn can lead to decreased leakage of the drug from the liposomes [43]. On the other hand, the results illustrated in **Figure 1a)** show that the presence of 30 mol% chol decreased the EE%. The highest EE% above 91% was obtained with the L/D of 30:1 and 60:1 without chol and in order to keep a relevant therapeutic dose, the 30:1 formulation was used for all further experiments. These results are congruent with other studies, where RAPA [45] and other hydrophobic drugs were encapsulated in chol-containing liposomes [46]. The accepted explanation for this is that the addition of chol in the phospholipid bilayer reduces the size of hydrophobic cavities, which leaves less space for the encapsulation of lipophilic drugs, decreasing the drug-lipid interactions and hence, the EE% [44–47]. **Table 1** shows the difference in size for loaded and empty liposomes measured with DLS in ultrapure water, where the size of RAPA-loaded liposomes slightly increased. Both systems had a low polydispersity index (PDI) of below 0.2. **Figure 1b)** and **c)** show that the liposomes were stable for a period of at least 16 weeks, with a slight decrease in size at week 2, which was already reported for DSPG/PC liposomes previously [32] and subsequent return to normal size at week 12.

**Table 1:**
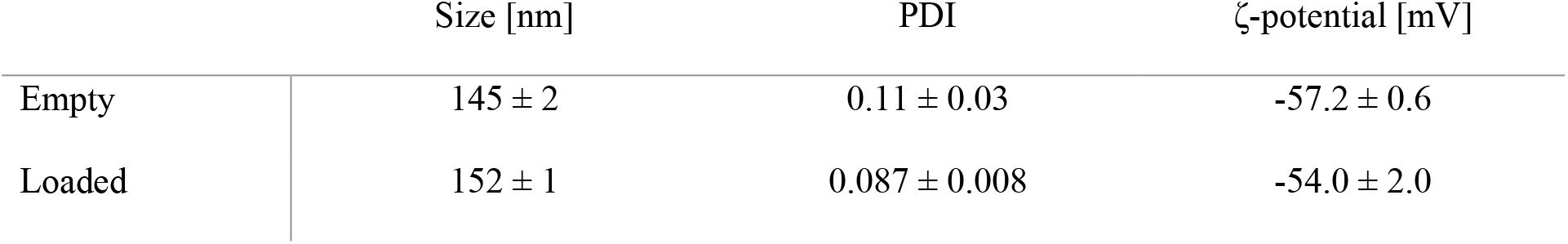
Liposomes’ properties at 30:1 L/D, without chol.

### 3.2 Fabrication and characterisation of zinc aggregated liposomes (ZnALs)

Small particles like liposomes are rapidly cleared from synovium and therefore the administration of larger particles is preferable [29]. Cationic vehicles have been recently used to enhance interaction with negatively charged cartilage surface [48,49], but liposomes containing high amounts of positively charged phospholipids can elicit toxicity and inflammation [50,51]. To avoid this, we devised a liposomal formulation with anionic phospholipids that aggregate after the addition of Zn^2+^, resulting in slightly positively charged ZnALs in μm-range (**Figures 2b, c)**.

**Figure 2:**
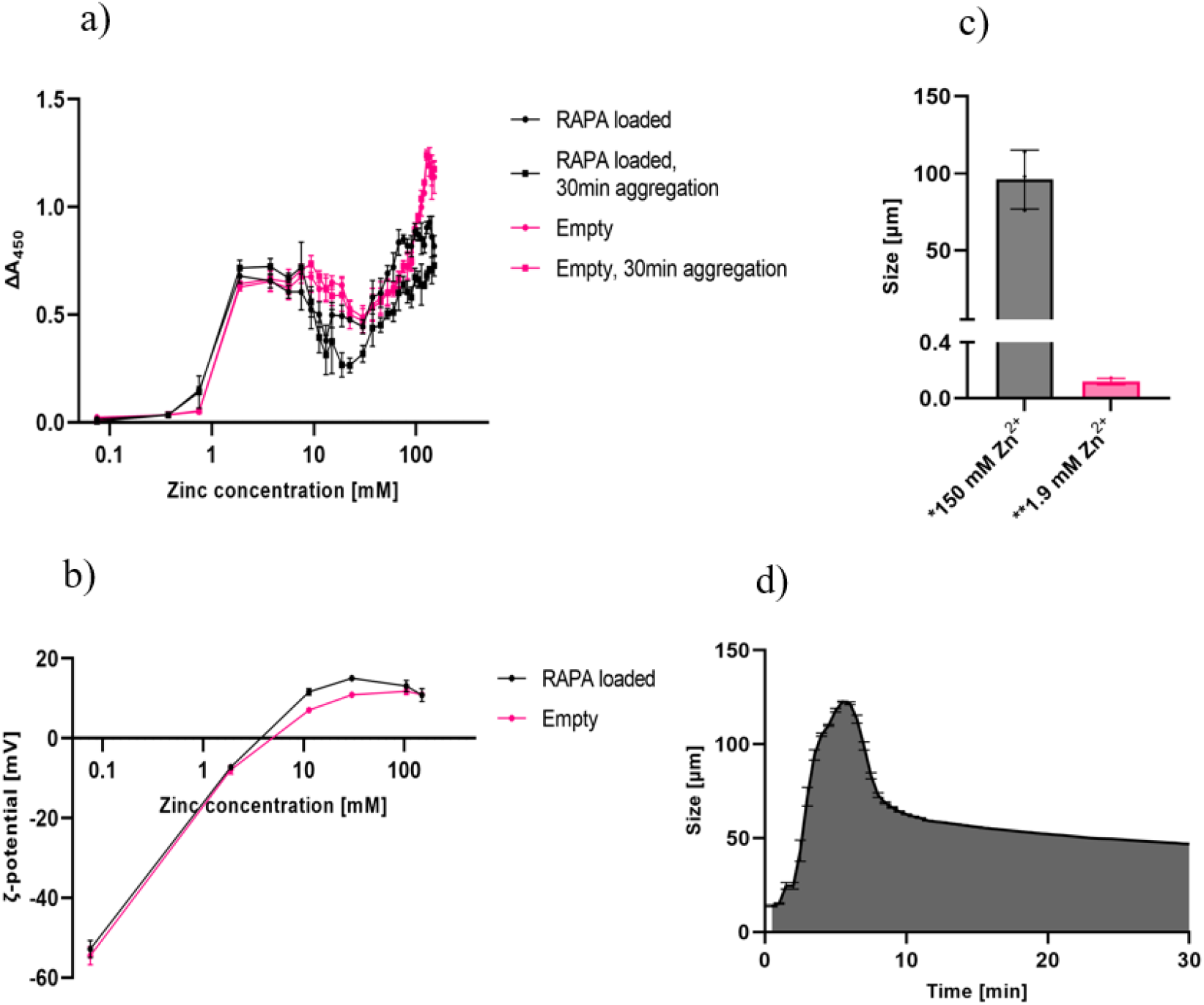
a) ZnALs’ aggregation profile measured at different Zn^2+^ concentrations via a plate reader at 450 nm, immediately after mixing or after 30 min of gentle mixing. Final lipid concentration was 5 mM b) ζ.-potential of ZnALs at different Zn^2+^ concentrations. c) Size after dilution with ultrapure water to a final Zn^2+^ concentration below 0.5 mM. *Measured with PSA. **Measured with DLS. d) Aggregation kinetics of liposomes in 150 mM Zn^2+^ measured with PSA – measurements were performed in 3 technical replicates at 30 s timepoints.

The aggregation is propagated by the neutralization of negatively charged liposomes with Zn^2+^, where the attractive van der Waals forces gain the upper hand. This is reflected in the secondary maximum on the aggregation profile on **Figure 2a)**. After further addition of the cation, more Zn^2+^ is bound on the surface of liposomes, which reverses the charge to +15 mV at 20 mM Zn^2+^ content and stabilises smaller particles. This phenomenon is seen as the secondary minimum in **Figure 2a)** and was previously observed in reports that focused on the coating of the anionic liposomes with polycations e.g., polylysine [52,53]. The continued introduction of Zn^2+^ to the final concentration of 150 mM induces a global maximum in the profile, where a dip in the ζ-potential to 11 mV can be observed for both empty and RAPA-loaded systems. The two systems exhibit distinct differences, with the former showing a less pronounced minimum at 20 mM Zn^2+^. This is likely due to a lower increase of ζ-potential at this concentration of Zn^2+^ to +11 mV compared to 15 mV for loaded ZnALs, which leads to a lesser stabilization of smaller particles. These shifts are presumably a consequence of RAPA’s perturbation of the phospholipid bilayer. The RAPA’s impact on the surface of DPPC-based anionic liposomes has been only recently elucidated [54], while the effect on the lipophilic moiety was studied earlier [55]. **Table 2** shows the results of the differential scanning calorimetry (DSC) analysis, where the impact of RAPA on DPPC/DSPG liposomal bilayer was evaluated on unextruded liposomes (MLVs) and aggregated vesicles (ZnALs). High rigidity of the used unsaturated phospholipids limited the impact of the encapsulated drug on the bilayer’s packing, as only a slight decrease in Tm can be observed. A more significant effect is the widening of the transition peak, suggesting that the homogeneity of the bilayers is affected by RAPA. Expectedly, the pretransition peak at 36 °C disappeared in the drug-loaded sample (**Table 2**), which indicates an interaction of the drug with the polar headgroups or acyl chains of the phospholipids. A greater effect of RAPA is observed in ZnALs, where both the transition and the pretransition peaks disappeared in the RAPA-loaded system, suggesting an altered packing of phospholipids. In summary, these DSC results show that RAPA affects the liposomal membrane, which could explain the differences in the aggregation behaviour in **Figure 2a)** and **b)**.

**Table 2:** DSC results of empty and RAPA-loaded MLVs and ZnALs (150 mM Zn^2+^). The change in enthalpy was normalized to the mass of phospholipids in the samples, while the reported results represent the calculated mean values ± standard deviation of at least 3 replicates. Thermograms are available in **Figure S2** in Supplementary Information.

| | $T_m$ [°C] | $\Delta H$ [J/g] | Peak width [°C] | T(pretr.) [°C] | $\Delta T$(pretr.) [J/g] |
| --- | --- | --- | --- | --- | --- |
| Empty MLVs | 43.66±0.03 | 35±3 | 3.27±0.03 | 36.0±0.2 | 2.0±0.4 |
| Loaded MLVs | 43.11±0.02 | 33±5 | 4.6±0.2 | n.a. | n.a. |
| Empty ZnAL | 43.9±0.1 | 1.0 ±0.6 | 2.3±0.2 | n.a. | n.a. |
| Loaded ZnAL | n.a. | n.a. | n.a. | n.a. | n.a. |

Subsequently, the size and reversibility of ZnALs were determined by dilution with ultrapure water to below the minimal concentration of Zn^2+^ that is needed for the aggregation (<0.5 mM). The measurements were performed with dynamic light scattering (DLS) and particle size analyser (PSA) that is based on laser diffraction technology, which is used to analyse particles in μm-range. The PSA measurements showed that the size distribution after 30 min of aggregation in 150 mM Zn^2+^ and subsequent dilution was unimodal with the mean size of 96 μm ± 19 μm (**Table 3** and **Figure 2c**)), indicating that the aggregates are irreversible. Upon performing the same procedure with the ZnALs at 1.9 mM Zn^2+^, the detector obscuration was found below 0.5%, which points to de-aggregation of ZnALs. Subsequent DLS measurement of the diluted samples revealed the reversion of ZnALs to their single-liposome size. However, for ZnALs at 150 mM Zn^2+^, the correlation function was irregular, and the size of aggregates could not be measured.

**Table 3:** Results of size measurements of ZnALs at different Zn^2+^ concentrations with DLS and PSA reported as mean ± standard deviation of at least 3 replicates.

| $[\text{Zn}^{2+}]$ in ZnALs | Size after dilution – DLS | Size after dilution – PSA |
| --- | --- | --- |
| 1.9 mM | 119 nm ± 20 nm | Not measurable |
| 150 mM | Not measurable | 96 $\mu\text{m}$ ± 19 $\mu\text{m}$ |

To confirm the presence of irreversible aggregates at 150 mM Zn^2+^, a nanoparticle tracking analysis (NTA) was used to capture the ZnALs on video after they were diluted 400x in ultrapure water and can be seen in **Supplementary Video 1**. In contrast, no aggregates can be observed in **Supplementary Video 2**, where ZnALs with 1.9 mM Zn^2+^ were diluted by the same dilution factor. The formation of irreversible aggregates was further studied by the addition of liposomes in excess volume of 150 mM Zn^2+^ solution, while the liquid was continuously flowing through a flow cell that was in a closed loop of PSA instrument. **Figure 2d)** shows the kinetic profile of size change for the period of 30 min. The aggregation reached its peak at 6 min of flowing through the flow cell at 122 μm and was followed by a decrease in size, which stabilised at 10 min. The gradual decrease in size over the following 20 min is likely due to shear stress, caused by the flow through the flow-cell. The morphology of liposomes and ZnALs were assessed with Cryo-TEM and the representative images are shown in **Figure 3**. The liposomes were mainly unilamellar and nanosized, consistent with the DLS data. The empty liposomes occasionally flattened, wheras the addition of 1.9 mM Zn^2+^, surprisingly, formed polyhedral structures with a negative curvature of the bilayer. Loaded liposomes only showed slight flattening at 1.9 mM and clear curvature at 150 mM Zn^2+^ (**Figure 3** and **Supplementary Video 3**).

**Figure 3:**
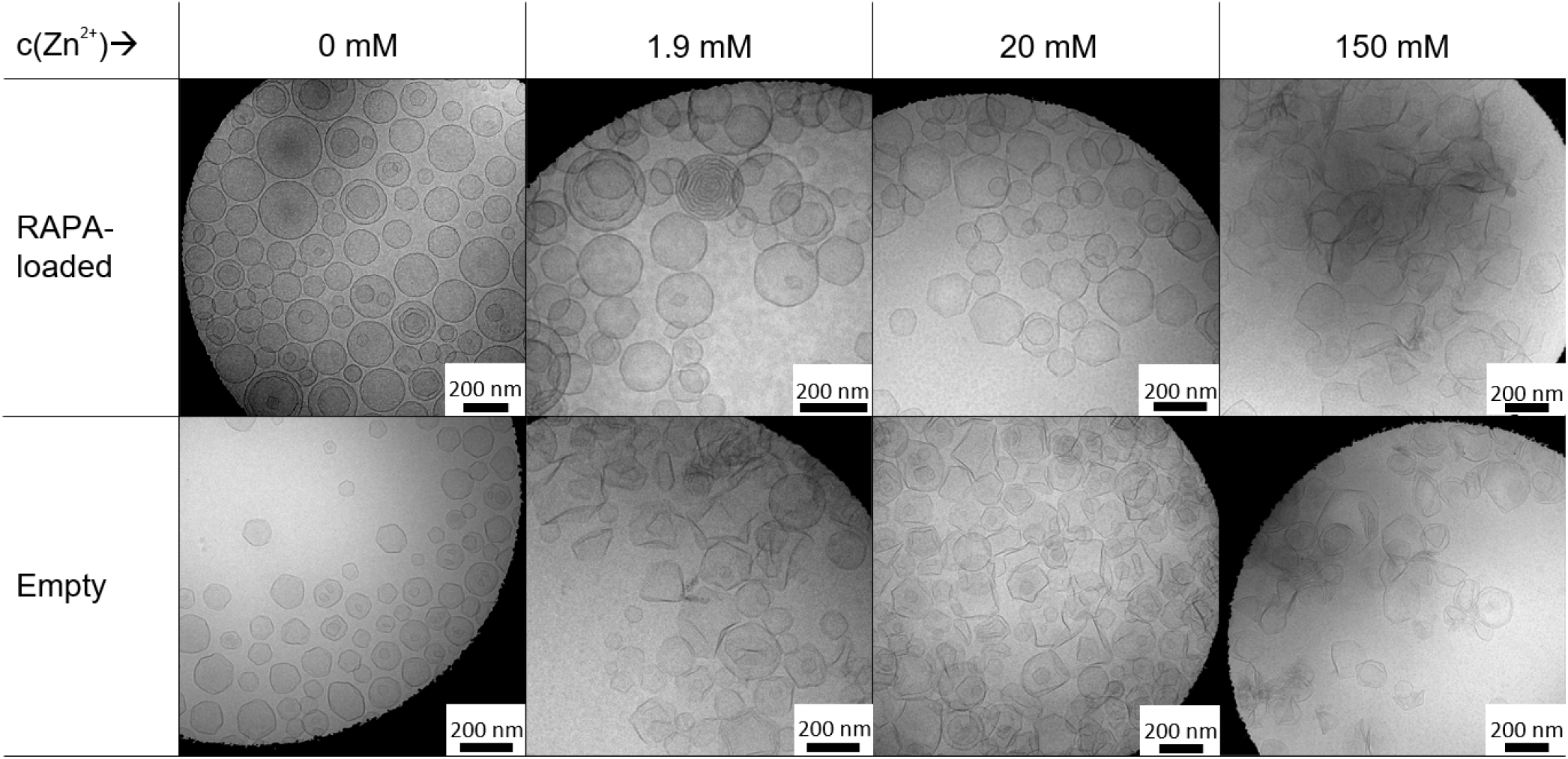
Cryo-TEM images of empty and RAPA-loaded formulations with different Zn^2+^ concentrations.

This was not seen in our previous study with DPPC/DSPG/chol liposomes exposed to 10 mM divalent cation, where only aggregation was observed [33]. The obvious explanation for this is that the particles in the present study have an enhanced fluid character due to the lack of chol in the bilayer and the formation of polyhedral structures is energetically favourable under the osmotic pressure imposed by external Zn^2+^. It has been previously reported that nanosized rigid liposomes can exist in a facetted configuration below their Tm [56]. The fact that RAPA-loaded liposomes exhibit a reduced angularity in up to 20 mM Zn^2+^ concentration suggests that the encapsulation of the drug in the bilayer has a chol-like effect on the membrane. This increased fluidity of the membrane is also supported by the DSC results in **Table 2**. The stabilization of liposomes’ spherical morphology by hydrophobic molecules is further supported by earlier investigation of the temoporfin encapsulation, which has comparable solubility in water to RAPA and has similarly reduced the angularity of the particles [57]. These results not only have importance for the design of liposomal drug delivery systems, but have further implications for better understanding of the cell membrane dynamics in different tonicities, as living cells also contain hydrophobic solutes. Furthermore, cryoTEM (**Figure 3**) confirmed the presence of aggregates at 1.9 mM and 150 mM Zn^2+^, which corroborated the PSA results.

Empty liposomes had clear aggregation at 20 mM Zn^2+^, while RAPA-loaded samples showed less pronounced aggregation as seen in **Figure 2a)**. Large aggregates were also scanned three-dimensionally using cryo-TEM tomography and are presented in **Supplementary Video 3**. To observe the system at a larger scale, liposomes were fluorescently labelled with DiD and aggregated as before (*vide supra*). **Figure S3** shows complex morphology of large ZnALs at 150 mM Zn^2+^.

### 3.3 RAPA release from ZnALs and effect on human OASFs

RAPA release from liposomes and ZnALs was tested in 10% EtOH in water to ensure sink conditions and stability of the drug throughout the experiment. **Figure 4a)** shows that ZnALs could retain the drug for longer than free liposomes with 86% released after 7 days, while 90% of the cargo was released from free liposomes already after 3 days. The use of aggregated liposomes for sustained drug delivery was recently proposed by our group in a study, where bupivacaine’s release was retained *in vitro* by calcium aggregated liposomes and the bioavailability of the drug was increased *in vivo* [33].

**Figure 4:**
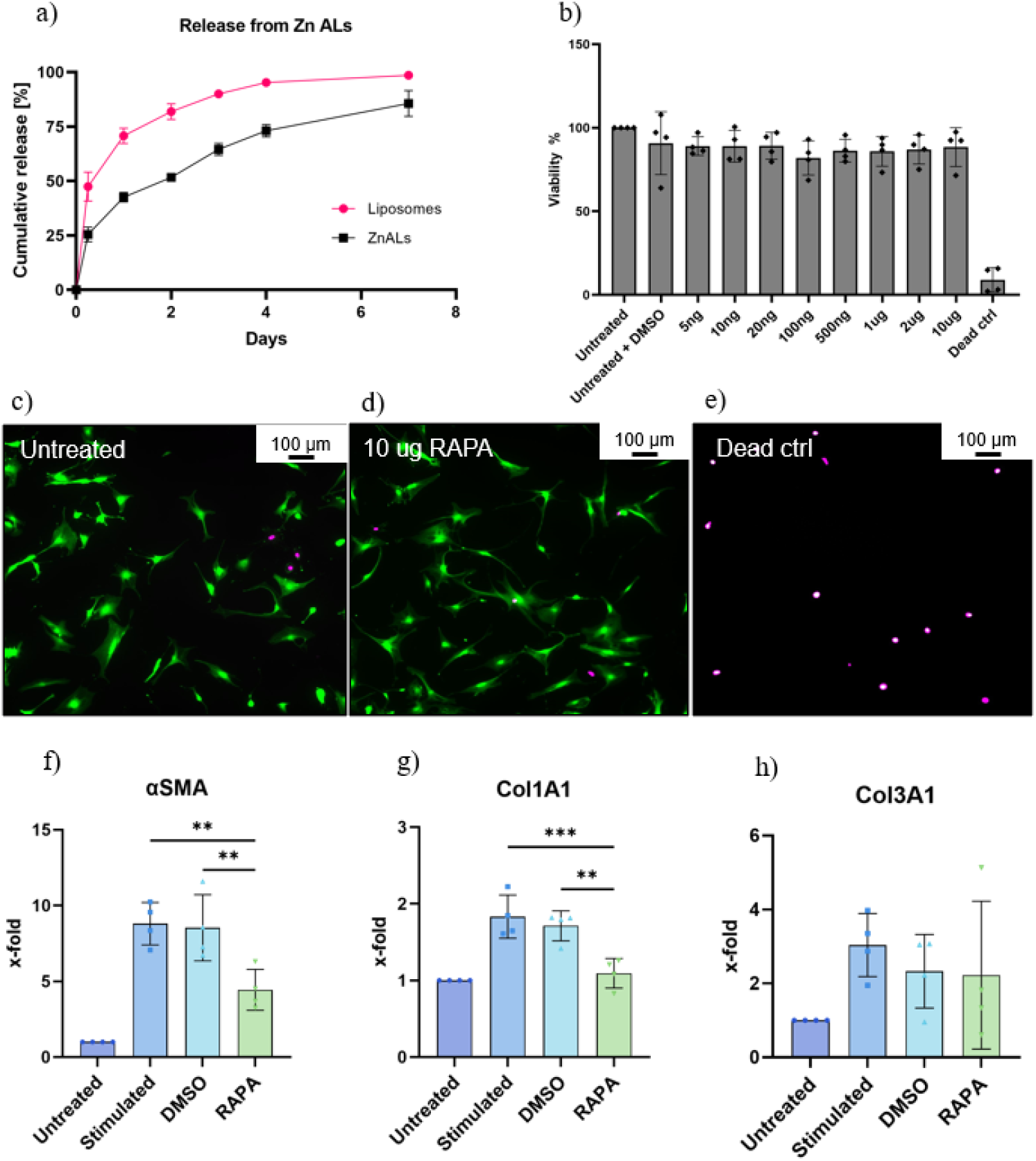
a) Release of RAPA from liposomes and ZnALs at 150 mM Zn^2+^ in 10% EtOH. b) Toxicity of RAPA on human OASFs stained with calcein; fluorescence was measured with a plate reader. c) Fluorescence microscopy image of untreated OASFs after 48 h incubation and subsequent staining with calcein and SYTOX Deep Red. Live cells are coloured in green and dead in magenta. d) Cells at 10 μg/mL RAPA. e) Dead cells treated with 70% EtOH. f) Gene expression of αSMA, g) Col1A1, h) and Col3A1 in OASFs that were stimulated with TGFβ (10 ng/mL) and treated with RAPA (1 μg/mL) for 48 h. One-way ANOVA and Tukey’s multiple comparisons test were run. Statistical significance is designated as: *P < 0.05 **P < 0.01, ***P < 0.001.

However, the previously reported aggregates needed an outside source of cations to retain the aggregated state and control the drug release. In our study, however, ZnALs were shown to be irreversible and exhibited a prolonged release even without the addition of Zn^2+^ in the release medium. These results suggest that ZnALs could potentially decrease the need for frequent intraarticular administration, which was previously connected with an increased risk of infections [28]. RAPA did not induce any toxicity in human OASFs after 48 h incubation with the drug in the range of 5 ng/mL up to 10 μg/mL, as reported in **Figure 4b-e)**. Next, the cells were stimulated with transforming growth factor-beta (TGFβ), a key mediator of synovial fibrosis in OA, to induce disease-like fibrotic response [2,16,17]. **Figure 4f)** and **g)** show that 1 μg/mL RAPA was able to decrease the gene expression of key profibrotic markers, in the OASFs, namely αSMA and Col1A1. Previous studies reported that RAPA can decrease inflammation and chondrocyte senescence in the synovial joints [14,58,59]. Our results add to this by showing that RAPA dampens fibrotic signaling in human OASFs *in vitro*. Furthermore, the toxicity of Zn^2+^ and RAPA in ZnALs was tested to find the dose for gene expression experiments. ZnALs were purified with dialysis prior to testing and the final Zn^2+^ concentration was measured with ICP-MS. RAPA content in purified ZnALs with 0.5 mM Zn^2+^ was 100 ng/mL and the ratio between the two was kept constant for higher doses (**Table S1**). No significant toxicity was observed at 0.5 mM of Zn^2+^, while 5 mM induced cell death (**Figure 5**).

**Figure 5:**
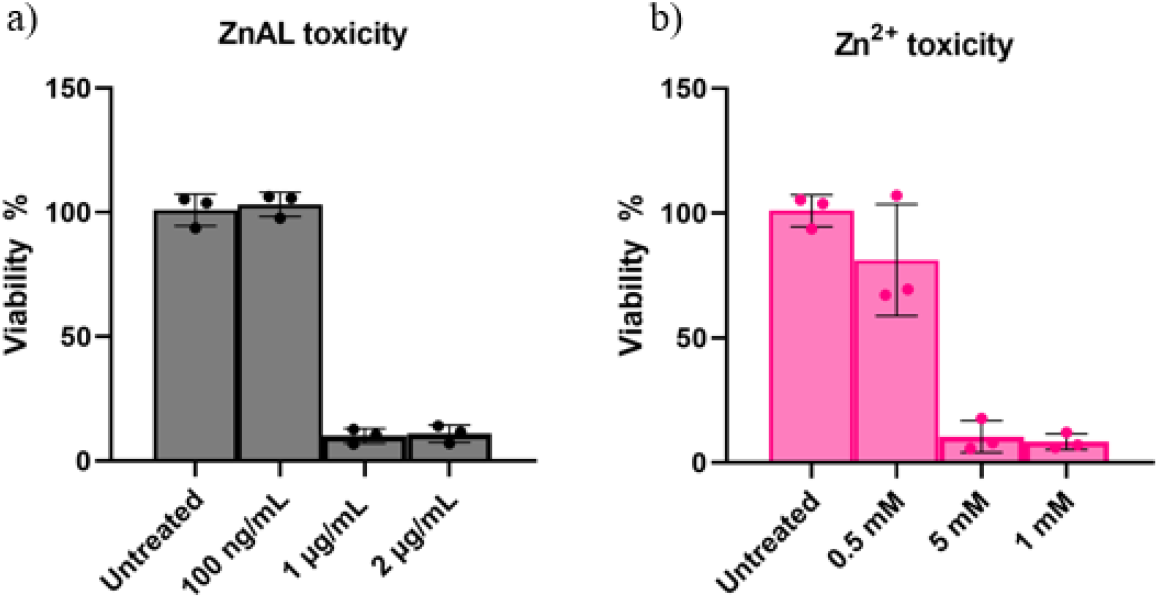
a) Toxicity of purified ZnALs with RAPA doses 100 ng/mL, 1 μg/mL, and 2 μg/mL. b) Toxicity of free zinc in concentrations that are present in purified ZnALs at above RAPA doses (**Table S1**).

The same zinc concentration is present in purified ZnALs with 1 μg/mL RAPA content (**Table S1**), where a similar toxic effect was observed. For these reasons, purified ZnALs with 100 ng/mL dose of RAPA were used for further cell experiments. The gene expression data in **(Figure 6a-c)** show that 100 ng/mL RAPA did not decrease fibrotic markers as effectively as 1 μg/mL. This calls for further research to find a more potent drug or a less toxic aggregating agent.

**Figure 6:**
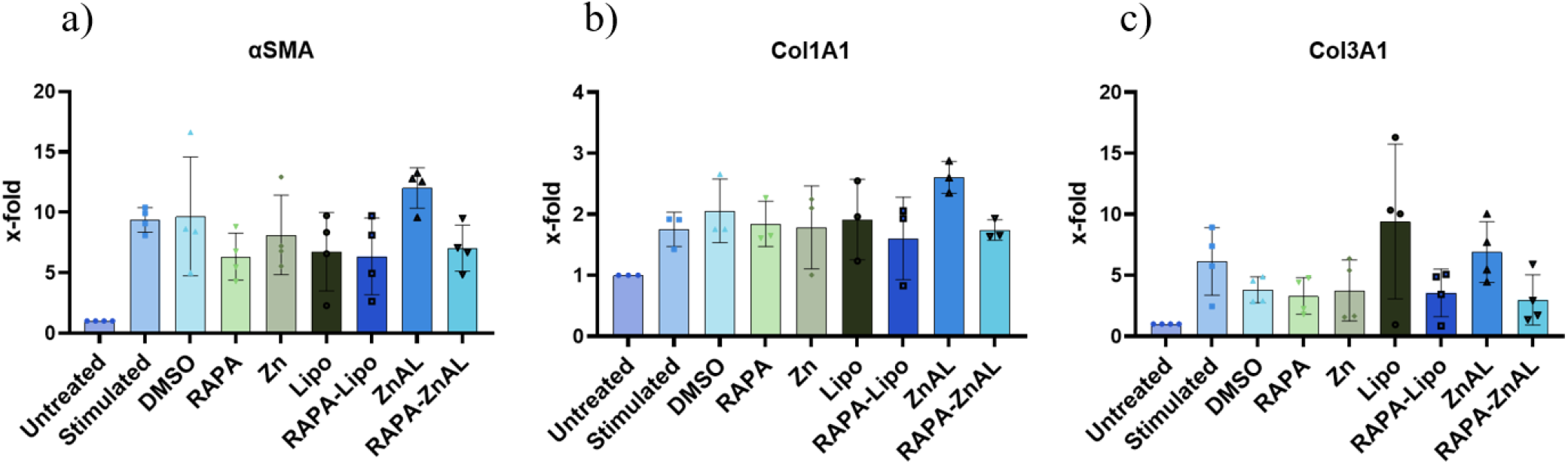
Gene expression of a) αSMA, b) Col1A1, n) and Col3A1 in OASFs that were stimulated with TGFβ (10 ng/mL) and treated with specified conditions for 48 h.

### 3.4 Lubrication of cartilage

Lubrication properties of the system were analysed with colloidal probe lateral force microscopy (CP-LFM) on silicon surface and a UMT-2 macro-tribometer with self-mated *ex vivo* porcine cartilage. Silicon surface that was used as a surface for CP-LFM measurement was cleaned with ethanol and treated with UV/ozone to assure cleanliness and a negative surface charge, which is present also in the *in vivo* cartilage conditions. **Figure 7a)** shows the change of friction force in dependence on applied normal force with the cantilever against the silicon surface that was treated with different conditions, as previously described [22]. The measured friction coefficient (COF) values obtained from the slope of the curve, drastically decreased when the surface was treated with liposomes and purified ZnALs (0.017) in comparison to PBS (0.279). It is generally accepted that the superior lubrication of liposomes stems from the formation of a hydration shell around the dipole of phospholipids, which can support high compressive loads and is very fluid [24,60]. A similar mechanism is expected here after the treatment with ZnALs, whose mean COF is lower compared to the liposomes’, which is 0.0279. This hypothesis is supported by the increase of friction after Zn^2+^ treatment, suggesting that no hydration layer is formed in the presence of Zn^2+^ only. This means that the lubrication observed with ZnALs is not due to the presence of the metal ions that increase the COF, but the presence of phospholipids.

**Figure 7:**
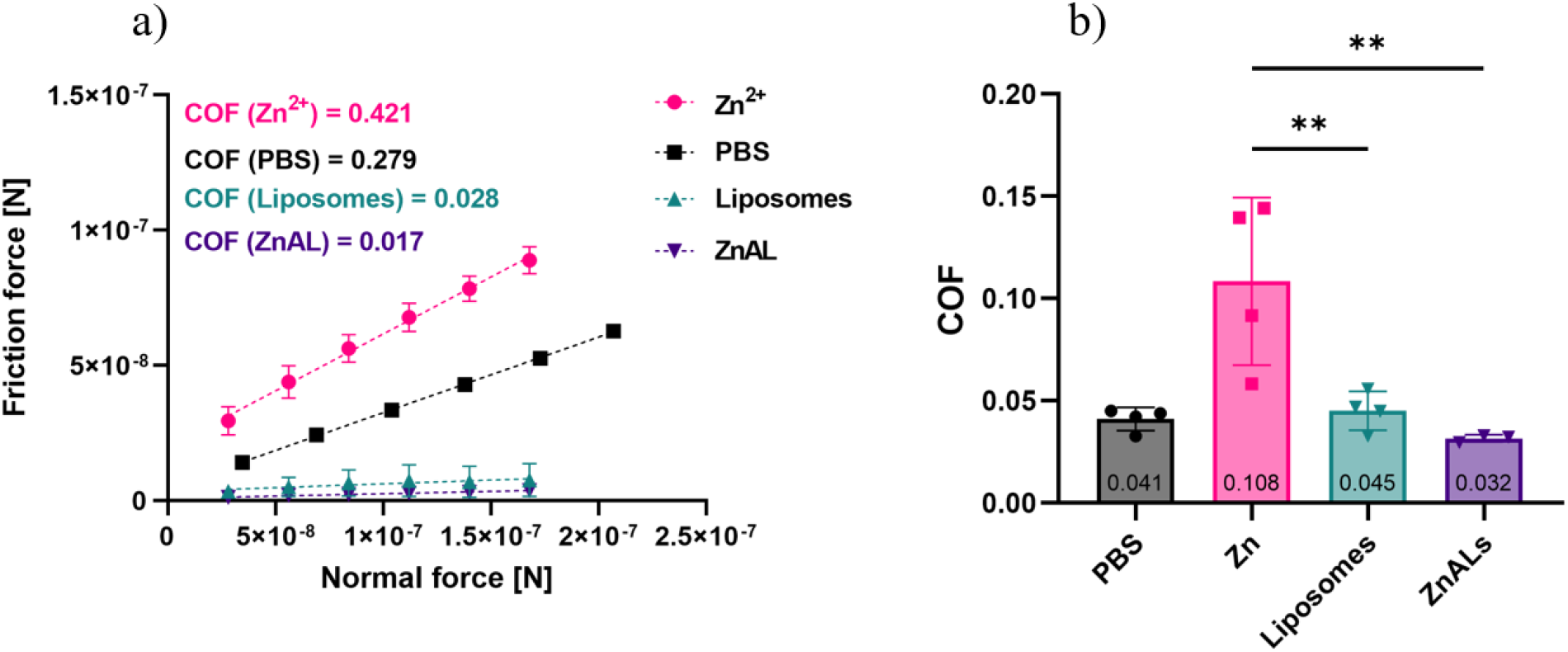
a) Friction as measured with CP-LFM on silica surface. b) Friction as measured with UMT on ex vivo porcine cartilage. ZnALs were purified before testing as described above. One-way ANOVA with Tukey’s multiple comparisons test was run. Statistical significance is designated as: *P < 0.05 **P < 0.01, ***P < 0.001, ****P < 0.0001.

A similar trend can be observed **Figure 7b)**, where the COF was measured on *ex vivo* cartilage with a macro-tribometer. The presence of zinc drastically increased the friction on the cartilage to a COF of 0.108 compared to the PBS of 0.041. This increase by the metal cations was negated by the electrostatic adsorption on the negatively charged liposomes and the formation of ZnALs, which significantly decreased the COF to 0.032. The mean COF of the free liposomes was measured at 0.045. Taken together, the formation of aggregates improved the friction both on nano- and macro-tribological scales, which indicates the potential of the formulation for the treatment of damaged cartilage in OA. More research is needed to confirm the observed responses under physiological conditions.

## 4 Conclusion

In the present report, we show that the aggregation of negatively charged liposomes with 150 mM Zn^2+^ yields irreversible aggregates (ZnALs) with a diameter above 90 μm, which was reported to drastically increase retention in synovial joints. We characterised the aggregation properties in depth and showed that ZnALs can sustain the release of RAPA, which decreases fibrotic markers in human OASFs. While necessary for the aggregation and the controlled release, Zn^2+^’s toxicity limits the therapeutic window of ZnALs. Aggregate formation prevents the increase of friction due to the presence of Zn^2+^ *ex vivo*, significantly decreasing the friction coefficient (COF) to 0.032, below the PBS control. The same effect was observed in a nanotribological setting, where ZnALs lowered the COF compared with free zinc and outperformed PBS. In summary, ZnALs are a drug delivery system that can sustain the release of RAPA longer than free liposomes, with 86 % of the drug released at 7-day mark. Future research should focus on improvements in the toxicity profile of the aggregated liposomes by finding new aggregating agents and thus improving the therapeutic effect of the formulation.

## 5 Acknowledgments

This work was supported by The Open Round grant by the Faculty of Science of the University of Bern. The authors thank the Center for Microscopy and Image Analysis at the University of Zurich for maintaining the imaging equipment and Dr. Elena Pachera for her support in acquiring images, as well as Peter Künzler and Benvinda Henriques Campos for their valuable support in the cell culture laboratories. Aggregation kinetics and aggregates’ size were measured and analysed with the support of Anton Paar TriTec SA in Neuchâtel. Fresh preparation of liposomes for cell experiments was possible thanks to the hospitality of Prof. Jean-Christoph Leroux and his team at ETH Zürich. The authors are grateful to Nicola Lüdi at Department of Chemistry, Biochemistry and Pharmaceutical Sciences of University of Bern for his support with ICP-MS measurements. The authors would also like to thank the workshop of Department of Chemistry, Biochemistry and Pharmaceutical Sciences of the University of Bern for making the dialysis devices for *in vitro* drug release experiments. Frank Steiniger of the University of Jena, Germany, is warmly acknowledged for his support with the cryo-TEM image acquisition and analysis of RAPA-loaded liposomes.

## 6 Author contributions

GB: Conceptualization; Funding acquisition, Investigation; Validation and Visualization; Formal analysis; Data curation; Writing of the original draft. SNR: Methodology (nanotribology); Investigation. SGE: Methodology (cell culture and gene expression). RE: Investigation; Validation. AA: Methodology (macrotribology). SA: Methodology. MFB: Conceptualization; Supervision. LI: Funding acquisition. RC: Funding acquisition; Supervision. OD: Funding acquisition; Supervision. PL: Conceptualization; Supervision; Project administration; Funding acquisition.

All authors contributed to the interpretation of the data and to drafting and revising the manuscript.

