## Supplementary Information for "Liposomal aggregates sustain the release of rapamycin and protect cartilage from friction"

S. N. Ramakrishna, L. Isa

[b] Department of Materials, ETH Zürich, HCl G 501, Vladimir- Prelog-Weg 1-5/10, 8093 Zürich, Switzerland

S. G. Edelat, O. Distler

[c] Center of Experimental Rheumatology, University Hospital Zürich, Wagistrasse 14, 8952 Schlieren, Switzerland

A. Arcifa, R. Crockett

[d] Laboratory for Surface Science and Coating Technologies, EMPA, Überlandstrasse 129, 8600 Dübendorf, Switzerland

M. F. Bertoncelj

[e] BioMed X Institute, Im Neuenheimer Feld 515, 69120 Heidelberg, Germany

1. Dialysis device for *in vitro* drug release

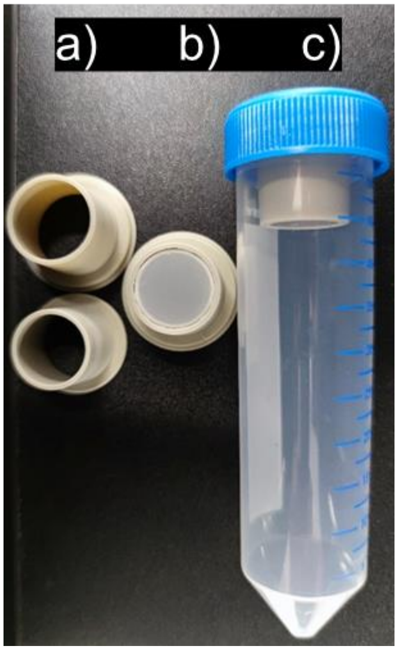


*Figure S1:* Dialysis device for drug release experiments a) dissasebled without a membrane, b) assembled with a 100 nm membrane and c) inside a 50 mL conical tube.

2. Primers for gene expression studies

αSMA Fwd: 5' GAC AAT GGC TCT GGG CTC TGT AA 3', Rev: 5'ATG CCA TGT TCT ATC GGG TAC TT 3'

Col1A1 Fwd: 5' CAG CCG CTT CAC CTA CAG C 3', Rev: 5' TTT TGT ATT CAA TCA CTG TCT TGC C 3'

Col3A1 Fwd: 5' GGA CCT CCT GGT GCT ATA GGT 3', Rev: 5' CGG GTC TAC CTG ATT CTC CAT 3')

RPLP0 Fwd: 5'-GCG TCC TCG TGG AAGTGA CAT CG 3', Rev: 5'-TCA GGG ATT GCC ACG CAG GG 3'

### 3. DSC thermograms

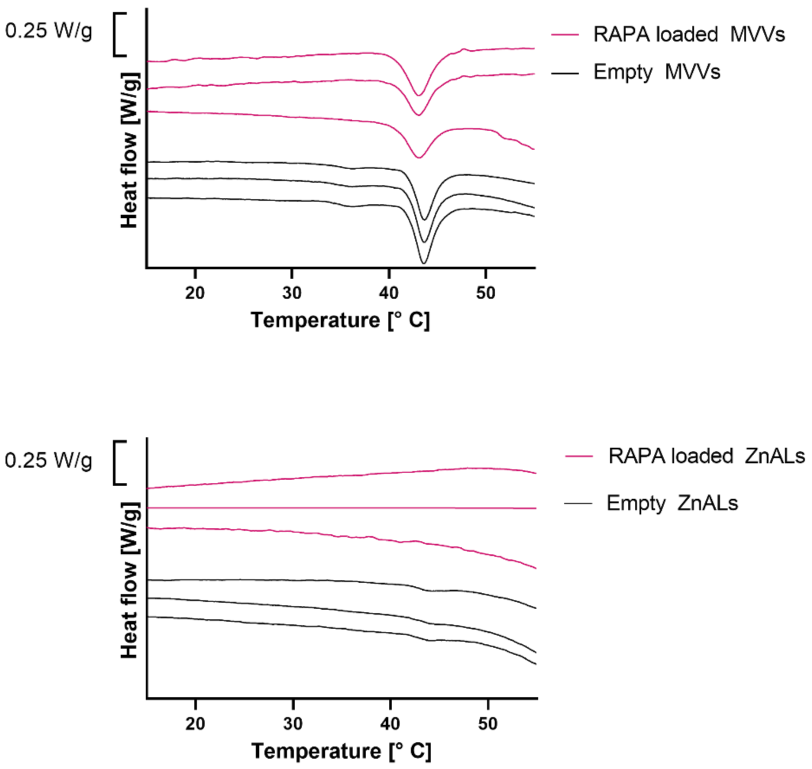


*Figure S2: DSC thermograms (heat flow endo down) of unextruded liposomes – multilamellar vesicles (MLVs) and of ZnALs with loaded RAPA and without (empty). Each thermogram is vertically shifted to avoid overlapping one another.*

### 4. Fluorescence microscopy images

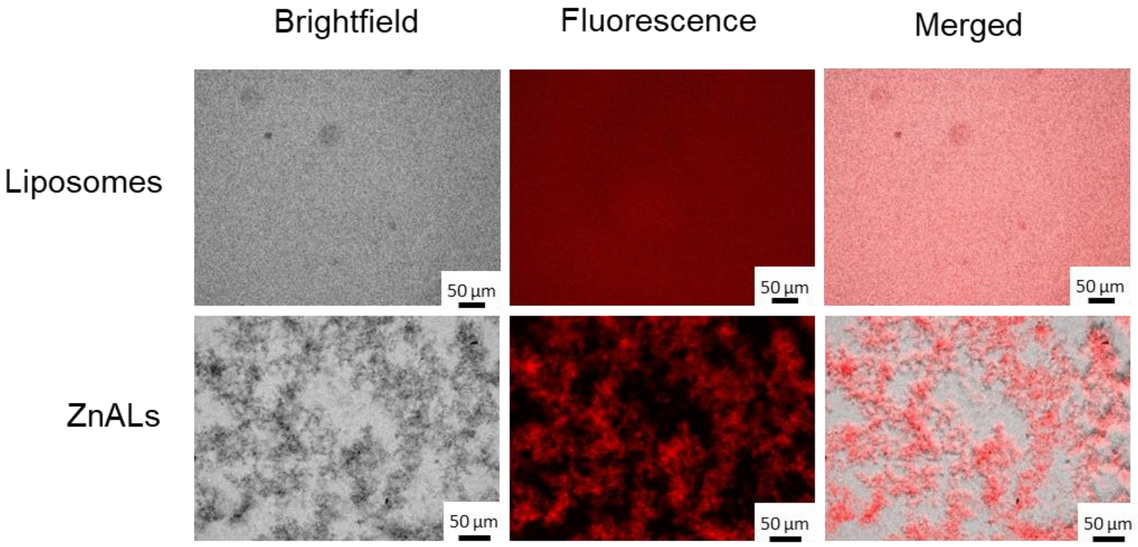


*Figure S3: Fluorescence microscopy images of liposomes and ZnALs at 150 mM of $Zn^{2+}$ and encapsulated fluorescent DiD probe.*

### 5. RAPA dose and Zn content in ZnALs

*Table S1: Dose of RAPA and $Zn^{2+}$ in ZnALs.*

| RAPA dose[$\mu$g/mL] | c($Zn^{2+}$) [mM] |
|---|---|
| 0.1 | 0.5 |
| 1 | 5.0 |
| 2 | 10 |
